# Sphingosine kinase 1 is induced by glucocorticoids in adipose derived stem cells and enhances glucocorticoid mediated signaling in adipose expansion

**DOI:** 10.1101/2024.09.13.612482

**Authors:** Johana Lambert, Anna Kovilakath, Maryam Jamil, Yolander Valentine, Andrea Anderson, David Montefusco, L. Ashley Cowart

## Abstract

Sphingosine kinase 1 (SphK1) plays a crucial role in regulating metabolic pathways within adipocytes and is elevated in the adipose tissue of obese mice. While previous studies have reported both pro- and inhibitory effects of SphK1 and its product, sphingosine-1-phosphate (S1P), on adipogenesis, the precise mechanisms remain unclear. This study explores the timing and downstream effects of SphK1/S1P expression and activation during *in vitro* adipogenesis. We demonstrate that the synthetic glucocorticoid dexamethasone robustly induces SphK1 expression, suggesting its involvement in glucocorticoid-dependent signaling during adipogenesis. Notably, the activation of C/EBPδ, a key gene in early adipogenesis and a target of glucocorticoids, is diminished in SphK1-/- adipose-derived stem cells (ADSCs). Furthermore, glucocorticoid administration promotes adipose tissue expansion via SphK1 in a depot-specific manner. Although adipose expansion still occurs in SphK1-/- mice, it is significantly reduced. These findings indicate that while SphK1 is not essential for adipogenesis, it enhances early gene activation, thereby facilitating adipose tissue expansion.

## Introduction

Sphingosine kinases, SphK1 and SphK2, phosphorylate sphingosine to form sphingosine-1-phosphate (S1P), a bioactive lipid in various cellular processes. The roles of SphK1 and S1P have been extensively studied in different tissues in the context of obesity and metabolic syndrome (1-3). In mice, constitutive deletion of SphK1 has been shown to protect against high-fat diet-induced insulin resistance, adipose inflammation, and liver steatosis (4). However, our recent studies demonstrate that adipocyte-specific deletion of SphK1 does not elicit similar protection and, in fact, exacerbates obesity-related pathologies (5). These findings suggest that SphK1/S1P can exert beneficial or deleterious effects depending on the cell type, tissue, and physiological context. This study focuses on elucidating the roles of SphK1/S1P in adipocyte precursor cells and their differentiation into mature adipocytes.

Adipose tissue is a dynamic organ that stores excess energy and regulates metabolism through the production and secretion of adipokines. Adipose expansion, which increases storage capacity and prevents lipotoxicity in peripheral tissues, occurs via adipocyte hypertrophy (increased cell size) and hyperplasia (increased cell number). Hypertrophy results from the accumulation of triglycerides in existing adipocytes, formed through lipogenesis, which includes *de novo* triglyceride synthesis from acetyl-CoA or re-esterification of free fatty acids with glycerol. Hyperplasia involves the proliferation and differentiation of adipose-derived stem cells (ADSCs) and pre-adipocytes into mature adipocytes, a process known as adipogenesis.

Subcutaneous adipose tissue more readily undergoes hyperplasia, whereas visceral adipose tissue favors hypertrophy. Hyperplasia is associated with more favorable metabolic outcomes, while hypertrophy is often accompanied by insulin resistance, inflammation, and hypoxia - hallmarks of adipocyte dysfunction (6-8).

Adipogenesis involves the differentiation of resident adipose tissue stem cells into lipid-laden adipocytes that express adipocyte-specific genes, such as fatty acid binding protein 4 (FABP4/AP2) and adiponectin (ADIPOQ). Mature adipocytes regulate energy balance through lipolysis and lipogenesis and produce adipokines (e.g., adiponectin, leptin, IL-6, TNF) that have various metabolic effects. Key regulators of adipogenesis include peroxisome proliferator-activated receptor gamma (PPARγ), the master regulator of adipogenesis, and CCAAT-enhancer-binding proteins (C/EBPs), which enhance adipogenic gene expression.

While the role of SphK1 in adipogenesis has been investigated by others, these studies have yielded conflicting findings. Some studies suggest that SphK1/S1P inhibit differentiation (9-12), by maintaining multipotency of ADSCs or promoting alternative fates, such as osteogenesis and chondrogenesis. Conversely, others indicate a pro-adipogenic role (13-16). These discrepancies may be due to differences in timing, dosage, and cell type. Indeed, it has been consistently demonstrated that S1P inhibits adipogenesis when added to cultured cells at supraphysiologic doses, affecting proliferation and viability (9, 11, 17). However, whether similar mechanisms apply to SphK1 and endogenous S1P is unclear. Taken together, these results illustrate a need to determine timing, conditions, and mechanisms by which SphK1/S1P signaling - via both intracellular and extracellular pathways - regulate adipogenesis.

## Materials and Methods

### Cell size analysis

Adipose sections were stained with hematoxylin and eosin (H&E) for adipocyte size analysis. Slides were imaged on a light microscope (Leica DMI1 microscope, Leica MC170 HD camera).

### Isolation of murine primary adipose derived stem cells (ADSCs)

All animal experiments conformed to the Guide for the Care and Use of Laboratory Animals and were in accordance with Public Health Service/National Institutes of Health guidelines for laboratory animal usage. Subcutaneous (inguinal and axial) fat pads were excised from 3-6-week-old male C57BL/6J mice (Jax, 000664), and rinsed in 1X phosphate-buffered saline with 1X antibiotic antimycotic solution (Millipore Sigma A5955). The tissue was transferred to digestion buffer (100 mM HEPES, 120 mM NaCl, 50 nM KCl, 5 mM glucose, 1 mM CaCl2, 0.1% collagenase, 1.5% bovine serum albumin) and minced into small pieces. Minced tissue was then transferred to a 50 mL conical tube with a 25 mL serological pipette. Tissue was incubated at 37ºC in a shaker at 20 x *g* for 30 minutes, with manual shaking and observation every 10 minutes until digested. The digest was filtered through a 100 µm cell strainer into a new 50 mL tube, diluted 2-fold with expansion medium (DMEM/F12 with 10% FBS and 1X antibiotic antimycotic) and centrifuged at 500 x *g* for 5 minutes. Floating lipid and media were aspirated, saving only the cell pellet. The pellet was re-suspended in expansion medium, filtered through a 40 µm cell strainer and plated onto a tissue culture flask. Media was changed after 2 hours to remove non-adherent cells and debris. ADSCs were maintained at 37ºC and 10% CO2 in expansion medium (changed every 2-3 days). Near confluency the cells were split and re-plated at 10,000 cells/cm^2^ on 6-well culture plates for adipogenesis experiments.

### Adipogenesis assay

Adipogenesis was induced 48 hours after cells reached confluency in adipogenic induction medium (DMEM/F12, 10% FBS, 1% penicillin/streptomycin, 10 µg/mL insulin, 1 µM dexamethasone, and 0.5 mM 3-Isobutyl-1-methylxanthine). On days 2 and 4 post-induction, the cells were fed with adipogenic maintenance medium (DMEM/F12, 10% FBS, 1% penicillin/streptomycin, 10 ug/mL insulin). Adipogenic culture medium (DMEM/F12, 10% FBS, 1% penicillin-streptomycin) was used on day 6 post-induction.

### Dexamethasone treatment

Dexamethasone (Sigma, D9402) was prepared in ethanol. Cells were treated in 1% fatty acid free BSA instead of FBS to prevent addition of exogenous S1P.

### RNA isolation and qPCR

Total RNA was isolated cells using Trizol (Invitrogen, 15596026) followed by RNeasy mini kit (Qiagen, 74106) extraction and column purification. cDNA was synthesized from 1 μg of total RNA using iScript Advanced cDNA Synthesis Kit (Bio-Rad, 1708890). Real time PCR was performed using a CFX96 Real-Time System (Bio-Rad) and SSoAdvanced Sybr (Bio-Rad, 1725272). Mean normalized expression was calculated by normalizing to the geometric mean of reference genes *Ppia* and *Tbp* (*i*.*e*., root2[Cq gene 1 ×Cq gene 2]) using the ^ΔΔ^Ct method. Mean normalized expression was calculated by normalizing to the expression of *Hmbs1* in liver tissue. Primer sequences are listed below:

**Table 1.**
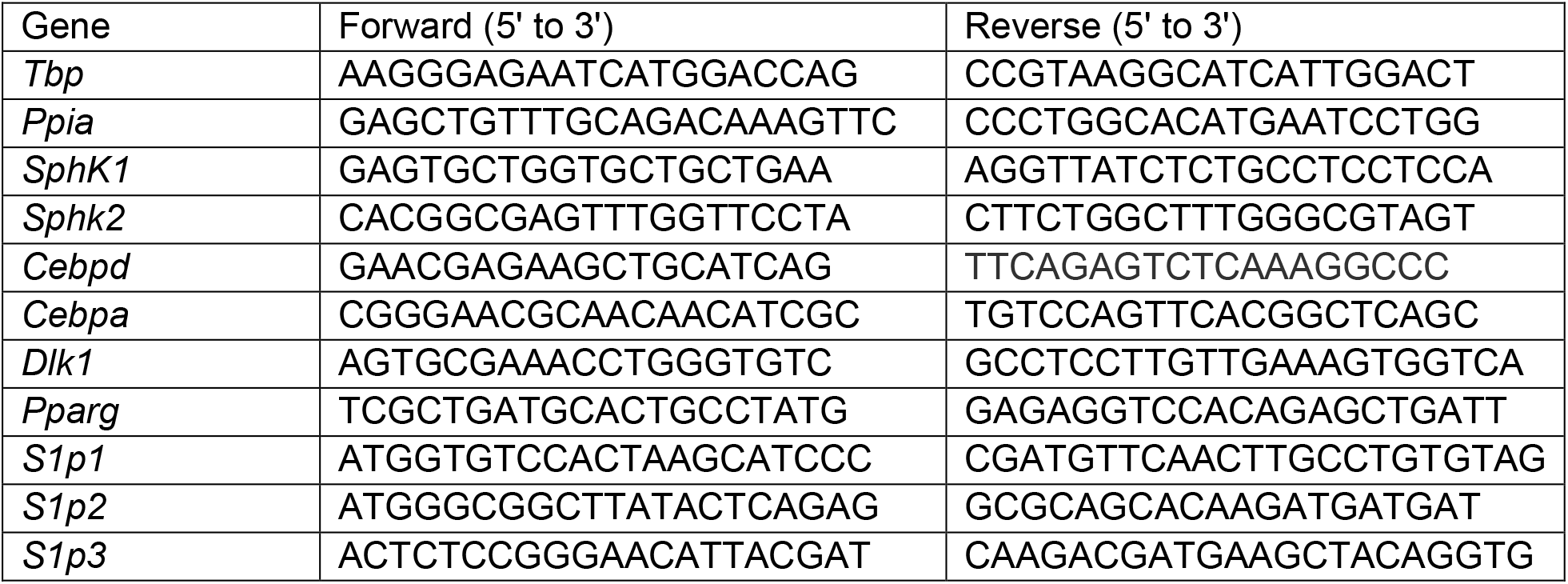
qPCR primer sequences.

### RNA sequencing

Gonadal adipose tissue RNAseq from SK1^fatKO^ mice and controls was conducted at the Medical University of South Carolina. Tissue was homogenized in Trizol followed by RNA isolation by RNEasy mini kit as described above. RNA integrity in tissue was assessed using the Agilent 2100 Bioanalyzer by the MUSC Proteogenomics Facility. All RNA samples had RIN > 7. 1 µg of RNA was submitted to the Genomics core facility at the Medical University of South Carolina and analyzed by the MUSC Bioinformatics Shared Resource.

### d17 sphingosine kinase assay

Cells were treated with 10 µM sphingosine (d17:1) (Avanti, 860640) for 30 minutes. Cells were rinsed with saline and collected using a cell scraper. Harvested cells were analyzed for d17 sphingosine and d17 S1P levels using liquid chromatography/tandem mass spectrometry (LC/MS/MS). d17 S1P measurements were normalized to total lipid phosphate.

### Oil Red O Staining

0.5% Oil Red O stock (Sigma, O-0625) was prepared in isopropanol by stirring overnight, then filtered to remove undissolved particles. Working solution was made just before using by mixing 6 parts Oil Red O stock with 4 parts diH2O for 20 minutes, then passed through a 0.2 µm filter. For staining, cells were fixed with 10% formalin for 1 hour. Wells were washed quickly with 60% isopropanol then allowed to dry completely. The Oil Red O working solution was added for 10 minutes and then rinsed with diH2O four times before imaging with a light microscope (Leica DMI1 microscope, Leica MC170 HD camera).

### Western blotting

Cells were rinsed twice in 1X PBS and harvested in RIPA buffer (150 mM sodium chloride, 50 mM tris-HCl, 1% Triton X-100, 0.5% sodium deoxycholate, 0.1% sodium dodecyl sulfate) with protease and phosphatase inhibitors. Cells were frozen and thawed to promote lysis, then centrifuged at 10,000 x *g* for 10 minutes to pellet cell debris. The supernatant was saved and protein content was determined using a bicinchonic acid assay (Thermo Fisher Scientific, 23225). Equal amounts of protein were separated by SDS-PAGE (Bio-Rad, Criterion TGX Stain-Free precast gels) and transferred to PVDF membranes. The membranes were blocked for 1 hour in 5% BSA. Primary antibodies for GR (Cell Signaling, 12041S) and phosphorylated GR (Ser211) (Invitrogen, PA5-17668) were diluted in 5% BSA. Proteins were detected using HRP-linked anti rabbit secondary (Cell Signaling Technology, 7074, 1:5000), Clarity ECL Western Blotting Substrate (Bio-Rad, 1705061) for HRP, and a ChemiDoc Imaging System (Bio-Rad, 17001401, 17001402). Vinculin (Cell Signaling Technology, 4650, 1:2000) and stain-free total protein were used to determine even loading. Band intensity was quantified using ImageJ.

### Corticosterone administration

Corticosterone was dissolved in ethanol. Mice were provided with 100 µg/mL corticosterone or 1% ethanol control in drinking water for 4 weeks starting at 14-16 weeks of age. The water bottles were replaced weekly and refilled as necessary. Mice were weighed weekly; at 4-weeks tissues were collected and snap frozen in liquid nitrogen.

### Glucose Tolerance Test

Intraperitoneal glucose tolerance tests were performed on mice after 3 weeks of corticosterone administration. Mice were fasted for 6 hours before conducting a glucose tolerance test, generally 08:00 hr to 14:00 hr. Mice received a sterile, intraperitoneal injection of 1.5 mg/kg D-glucose. Blood was collected through a nick in the tail and analyzed neat using a One Touch UltraSmart Blood Glucose Monitoring System at fasting for baseline blood glucose concentration and was then assessed 15, 30, 60, and 120 minutes after glucose injection.

## Results

To better understand how SphK1 deletion in adipocytes affects adipose tissue function, we conducted RNA sequencing on gonadal adipose tissue homogenates from SK1^fatKO^ mice and controls. The results revealed dysregulation of several key adipogenic genes: pro-adipogenic gene expression was reduced, while anti-adipogenic genes were upregulated in SK1^fatKO^ adipose tissue (Figure 1A). Our previous study also revealed a notable increase in adipocyte hypertrophy within the gonadal adipose tissue depot of SK1^fatKO^ mice (5). Subcutaneous adipose tissue, known for its susceptibility to expansion through adipogenesis more so than visceral adipose tissue, and is associated with glucose tolerance due to its ability to promote hyperplasia rather than hypertrophy. This type of expansion is critical to prevent inflammation, insulin resistance, and fibrosis both within adipose tissue and systemically (18). Given the importance of subcutaneous adipose tissue expansion in maintaining metabolic health, we examined adipocyte size in the inguinal (subcutaneous) depot of SK1^fatKO^ mice. Indeed, high-fat diet fed SK1^fatKO^ mice exhibited more than a 2-fold increase in average subcutaneous adipocyte size (Figure 1B), similar to the hypertrophy previously reported in the gonadal depot (5). These findings suggest that SphK1 deletion in adipocytes impairs adipogenesis, prompting further investigation into mechanisms by which SphK1 regulates adipogenesis.

**Figure 1:**
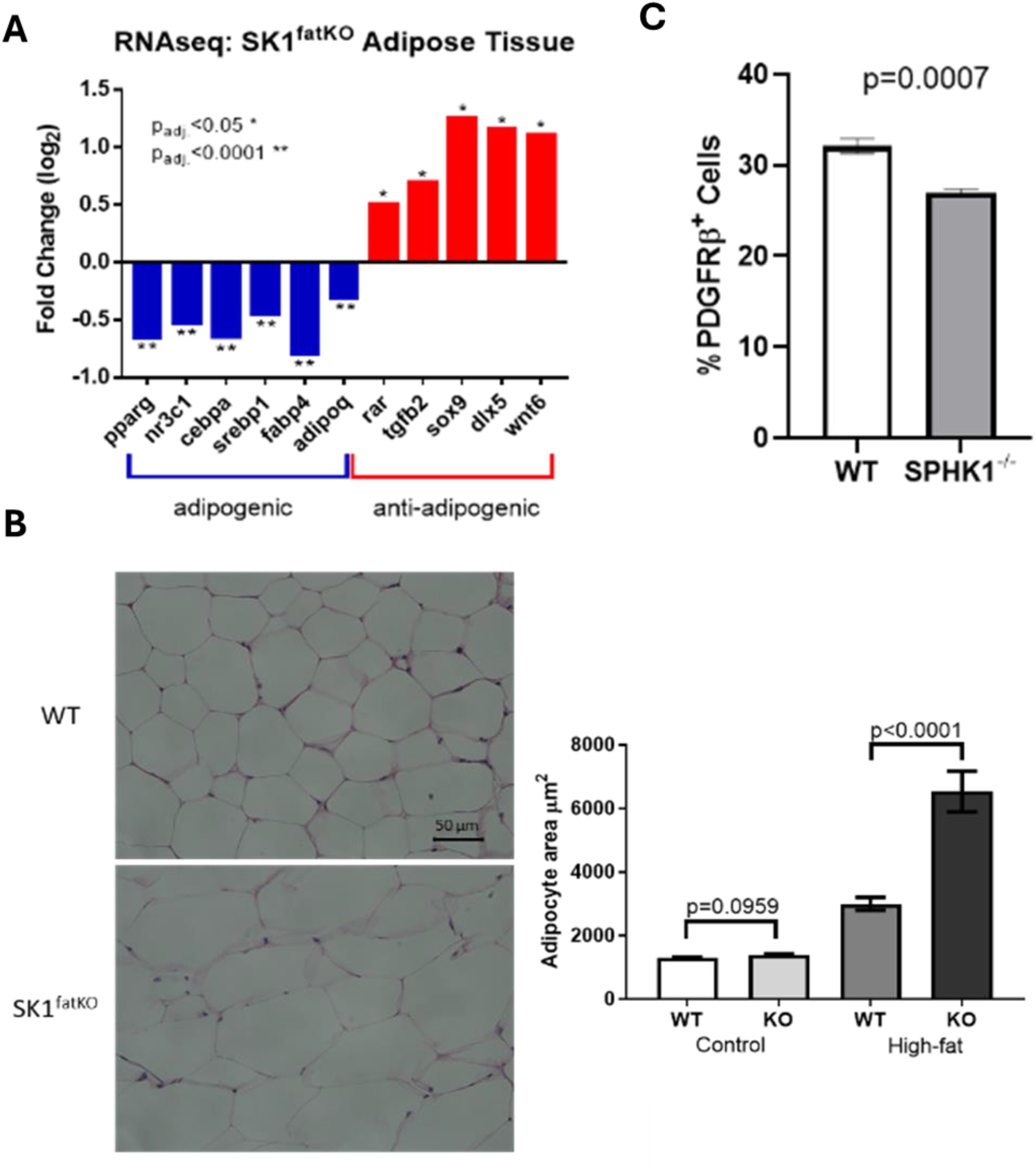
Hypertrophy and adipogenic markers in models of sphingosine kinase 1 deletion. **A**. Selected pro- and anti-adipogenic genes from RNAseq of SK1_fatKO_ adipose tissue. **B**. Size of subcutaneous adipocytes from adipose tissue of SK1_fatKO_ mice or controls on high-fat or control diet. **C**. Percent of PDGFRβ positive cells in the stromal vascular fraction of inguinal adipose tissue from SK1_-/-_ mice. Bars represent mean ± SEM; n = 3. *P < 0.05; **P < 0.0001. A. and C. unpaired t test. B. One way ANOVA.

Platelet-derived growth factor receptor β (PDGFRβ) is critical for vascular development and differentiation of many cell types. Expression of PDGFRβ in stromal vascular cells from adipose tissue is linked with high adipogenic capacity (19). While there is no definitive panel of markers to assess adipogenic potential or lineage commitment, PDGFRβ expression is an established marker that reliably predicts adipogenic potential *in vitro* (19-22). Thus, we isolated stromal vascular cells from inguinal adipose depot of constitutive knockout (SphK1^-/-^) mice and controls to determine if SphK1 depletion affects PDGFRβ expression in this cell population. The stromal vascular fraction is heterogenous, comprising ADSCs, committed preadipocytes, immune cells and endothelial cells. Significantly fewer PDGFRβ-positive cells were isolated from SphK1^-/-^ mice, suggesting a reduction in adipogenic commitment as a consequence of SphK1 depletion (Figure 1C).

The adipocyte hypertrophy and dysregulation of adipogenic genes in SK1^fatKO^ mice, along with reduced PDGFRβ expression in SphK1^-/-^ stromal vascular cells imply roles for SphK1 during both adipocyte commitment and maturation. Generation of the SK1^fatKO^ mouse utilized Cre recombinase driven by the adiponectin promoter, leading to adipocyte-specific gene deletion, since adiponectin is expressed only during later stages of adipocyte maturation. This model is useful for investigating SphK1 in mature adipocytes; however, as SphK1 is still expressed in earlier stages of adipogenesis, the effects observed may be due to cross-talk with existing mature adipocytes or changes within the adipose tissue microenvironment. To study the functions of endogenous SphK1 through all stages of adipogenesis – from adipocyte commitment to maturation - ADSCs obtained from constitutive SphK1^-/-^ mice offer a more appropriate *in vitro* system.

To begin investigating the actions of SphK1/S1P in preadipocyte commitment and differentiation, we assessed SphK1 expression and activity over an adipogenesis time course. ADSCs were isolated from the inguinal and axial (subcutaneous) adipose depots, expanded, and plated for differentiation. Following induction, *SphK1* expression peaked within 24 hours and then declined through the course of adipogenesis (Figure 2A). As SphK2 is an additional source of S1P, we also measured *Sphk2* mRNA over the time course, finding that it remained consistently lower than SphK1 and did not significantly change throughout differentiation (Figure 2B). Consistently, measurements of cellular S1P and SphK activity also indicated SphK activation within 24 hours of induction, followed by a decline by 48 hours (Figure 2C). Since SphK activity was minimal in SphK1^-/-^ ADSCs (data not shown), along with low SphK2 expression (Figure 2B), we primarily attributed changes in SphK activity during adipogenesis to SphK1 (Figure 2D).

**Figure 2:**
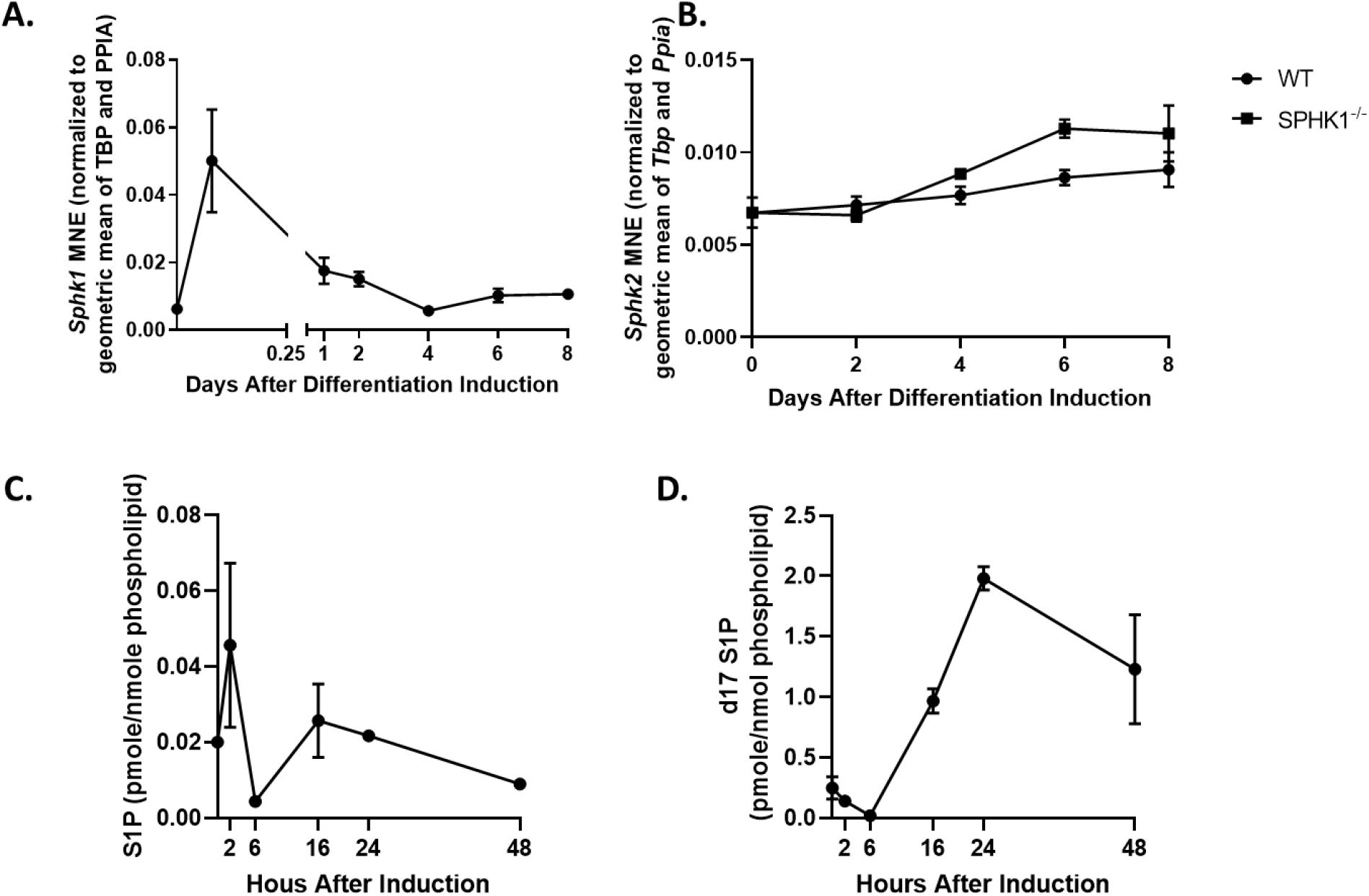
Measurements of SphK1 throughout adipogenesis. **A**. *SphK1* mRNA **B**. *Sphk2* mRNA **C**. Cellular S1P through early differentiation measured by mass spectrometry **D**. SphK activity. Each point represents mean ± SEM; n = 3.

During the first 24 hours of differentiation, significant changes in gene expression and cell morphology occur, driven by a cocktail of pro-adipogenic reagents. Standard *in vitro* adipogenesis protocols include insulin to stimulate transport of glucose and fatty acids for lipid synthesis; IBMX, to raise cellular cAMP; dexamethasone, a synthetic glucocorticoid that induces expression of key genes such as C/EBPs; and sometimes rosiglitazone, a PPARγ agonist. Confluent, growth-arrested ADSCs are treated with these reagents for two days to induce adipogenic commitment and are then maintained with insulin-containing media throughout differentiation to promote lipogenesis and adipocyte maturation. To test the effects of SphK1 deletion on adipogenesis, we performed adipogenesis assays using ADSCs from SphK1^-/-^ and control mice. Surprisingly, SphK1^-/-^ ADSCs differentiated into adipocytes similarly to controls, both with and without rosiglitazone. Microscopy and triglyceride measurements revealed an approximately 25% reduction in lipid accumulation in both SphK1^-/-^ cells and controls, but no significant differences were observed between the genotypes (Figure 3A-B). Interestingly, protein levels of key adipocyte genes PPARγ and FABP4 were lower in SphK1^-/-^ cells compared to controls during the first 2 hours after induction, but by 48 hours, expression of PPARγ and FABP4 in SphK1^-/-^ cells surpassed that of controls (Figure 3C). Taken together, these results suggest a dual role for S1P in adipogenesis, supporting early adipogenic processes while interfering with later stages of differentiation.

**Figure 3:**
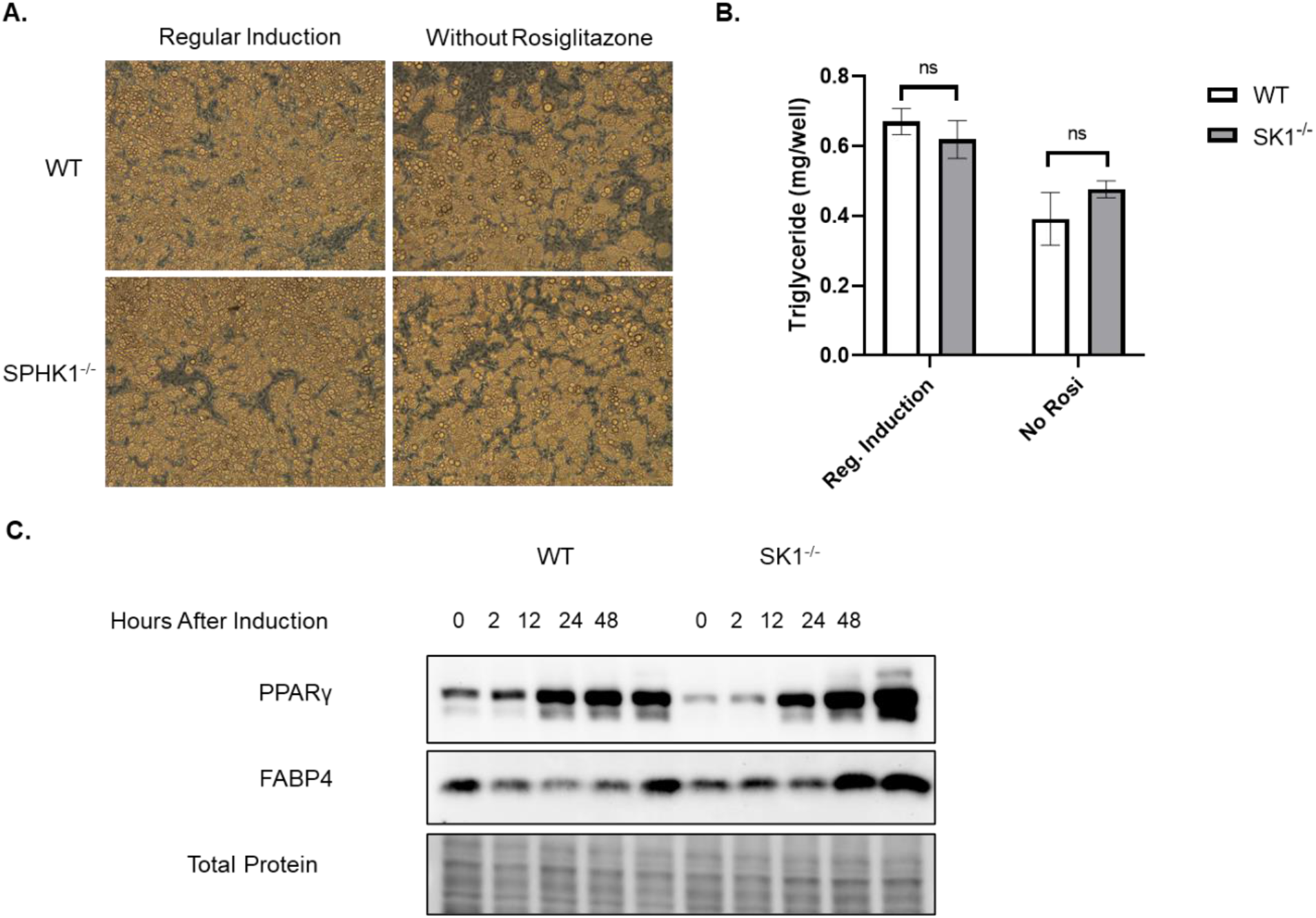
Adipogenesis assays and readouts. **A**. Images at 8 days post-adipogenic induction. **B**. Triglyceride measurements at 8 days post-adipogenic induction. **C**. Western blots for key adipocyte genes during early adipogenesis. Bars represent mean ± SEM; n = 3. **B**. unpaired t test. ns = not significant.

The typical adipogenic induction medium often employs a “sledgehammer approach” to induce adipogenesis in all cells simultaneously, using high doses of synthetic chemicals. This differentiation cocktail rapidly activates key adipogenic transcription factors, such as PPARγ and C/EBPα, potentially overriding mechanisms involving SphK1 that are relevant to *in vivo* processes. Additionally, fetal bovine serum (FBS) is used at 10% throughout the culture and adipogenic induction of ADSCs. Since blood components, including FBS, are known to contain high levels of S1P and other sphingolipids, we analyzed sphingolipid levels in FBS using mass spectrometry. Even when diluted to 10% in the media, S1P was found at nanomolar concentrations. Charcoal stripping of FBS reduced S1P levels by about half, though they remained within the range of S1P receptor KDs (23) (Figure 4).

**Figure 4:**
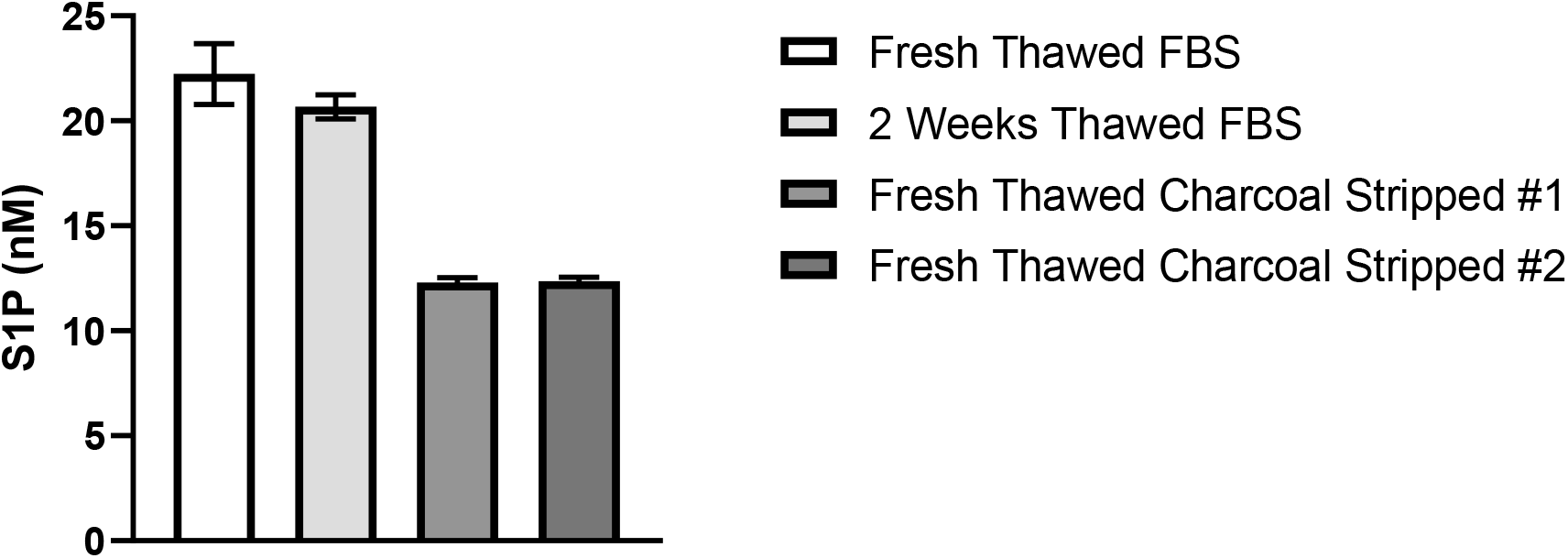
S1P concentration of regular or charcoal stripped 10% FBS. To investigate the cause and effects of SphK1 induction during *in vitro* adipogenesis, ADSCs were treated with individual pro-adipogenic media components (insulin, IBMX, dexamethasone, and rosiglitazone). This allowed us to examine the impact of each component on SphK1 expression while circumventing confounding effects from the full adipogenic cocktail or FBS. Interestingly, only dexamethasone induced a significant increase in SphK1 expression (∼5-fold), which was greater than the combined effects of the other reagents (∼2-fold) (Figure 5A). Sphk2 expression remained low and was not significantly altered by any of these reagents (Figure 5B).

To further characterize dexamethasone induction of SphK1, a dose response and time course were conducted. The dose response revealed that a concentration as low as 20 nM was sufficient to achieve maximal SphK1 induction (Figure 5C), although 1000 nM dexamethasone is typically used in adipogenesis assays. SphK1 induction occurred at the earliest time point measured after dexamethasone treatment (2 hours) and persisted for at least 72 hours (Figure 5D). Additionally, dexamethasone treatment stimulated SphK activity (measured by d17 sphingosine conversion to d17 S1P) at both 24 and 48 hours (Figure 5E). These results also confirm that SphK1 expression is indicative of SphK1 activity, a notable finding given the lack of reliable, commercially available antibodies for mouse SphK1. Interestingly, expression of S1P receptors 1-3 were all reduced after 24 hours of dexamethasone treatment (Figure 5F).

**Figure 5:**
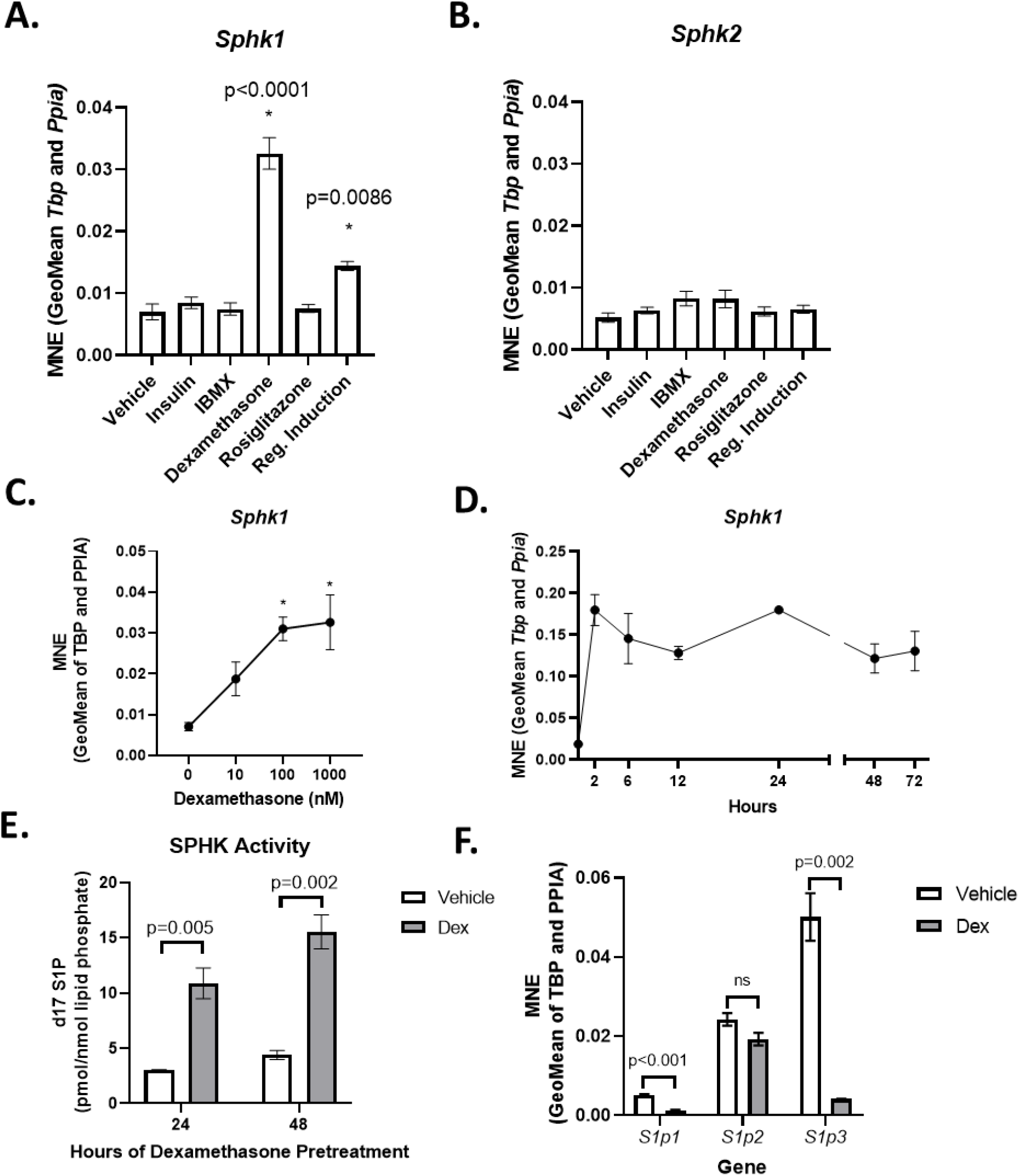
Measurements of SphK1 in ADSCs treated with pro-adipogenic media components or dexamethasone alone. A-B. *SphK1* and *Sphk2* expression in ADSCs treated with pro-adipogenic media components. **C**. *SphK1* expression in ADSCs treated with 0-1000 nM dexamethasone. **D**. *SphK1* expression in ADSCs treated with dexamethasone for 0-72 hours. **E**. SPHK activity in ADSCs treated with 100 nM dexamethasone. **F**. Expression of S1P receptors in ADSCs treated with 100 nM dexamethasone. Bars or points represent mean ± SEM; n = 3. *P < 0.01. A. and B. One way ANOVA. E. and F. unpaired t test. ns = not significant.

Given the limitations of *in vitro* adipogenesis assays, we investigated how dexamethasone alone affects adipogenic gene expression in SphK1^-/-^ compared to wild-type controls, this time in the absence of FBS. In cells treated with dexamethasone, a small but significant reduction of Cebpd activation, a key transcription factor in adipogenesis, and known glucocorticoid receptor target, was observed (Figure 6A). Activation of Cebpa, a downstream target of Cebpd in adipogenesis, was also decreased in SphK1^-/-^ ADSCs (Figure 6B). To determine if exogenous S1P could stimulate C/EBPδ, 150 nM S1P was added to ADSCs for 16 hours. No changes in Cebpd or Cebpa were observed (Figure 6C-D). S1P receptor agonists were also added throughout adipogenesis to test for potential pro-differentiation effects; however, no effects on differentiation were observed over a wide range of concentrations (data not shown). These experiments suggest SphK1 affects intracellular signaling during adipogenesis, while S1P receptor-mediated signaling is not a primary regulator of adipogenesis.

**Figure 6:**
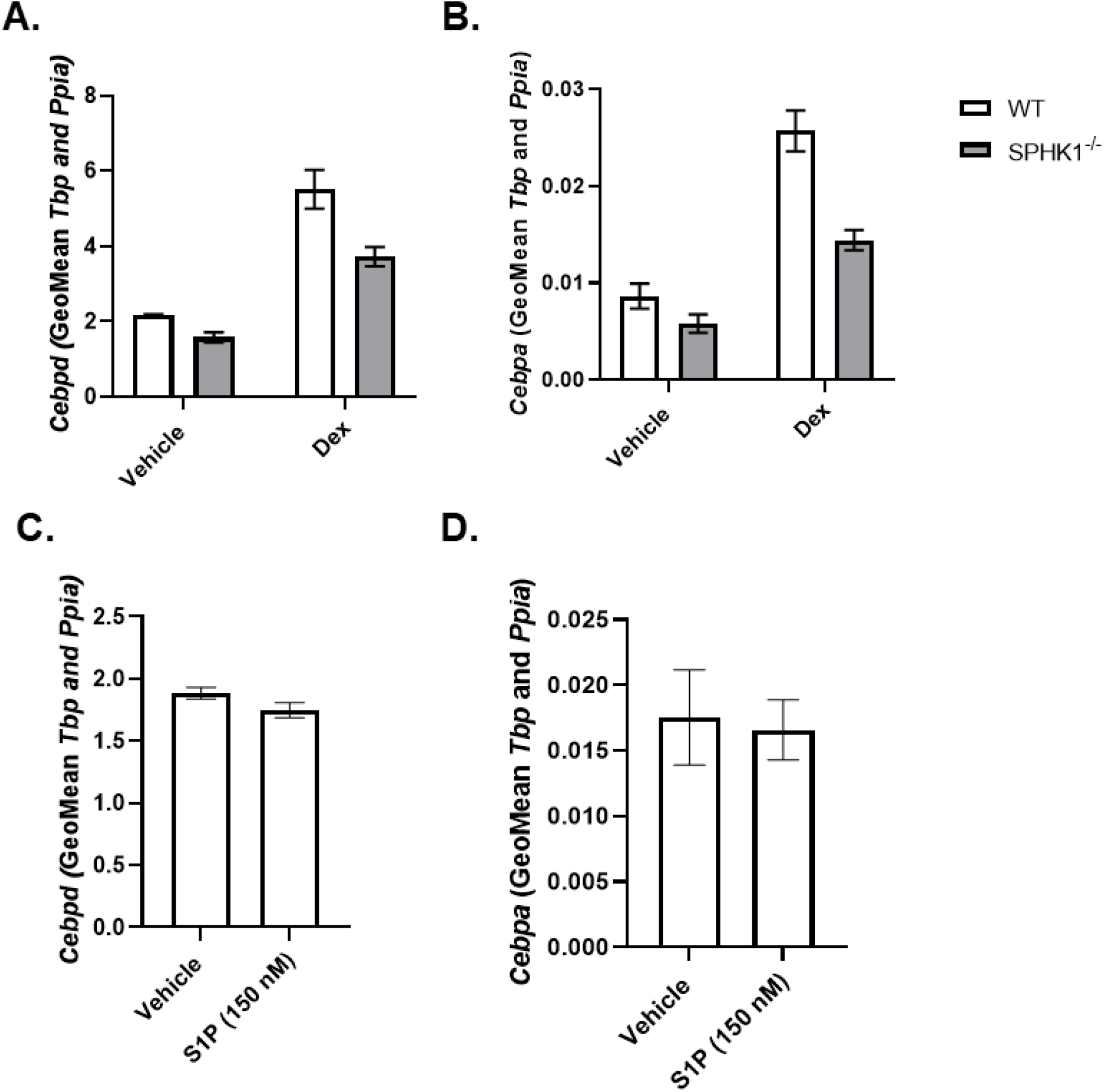
Dexamethasone stimulates SphK1-dependent C/EBP activation in ADSCs, which cannot be replicated by S1P treatment. **A-B**. *Cebpd and Cebpa* expression in dexamethasone-treated SphK1_-/-_ and control ADSCs. **C-D**. *Cebpd* and *Cebpa* expression in control ADSCs. Bars represent mean ± SEM; n = 3. A.-D. unpaired t test. ns = not significant.

To further elucidate the role of SphK1 in glucocorticoid signaling, we treated WT and SphK1^-/-^ cells with dexamethasone, then fractionated them to isolate cytosolic and nuclear proteins. Upon binding glucocorticoids such as dexamethasone, the glucocorticoid receptor (GR) dimerizes and becomes hyperphosphorylated before translocating to the nucleus to regulate gene expression by binding glucocorticoid response elements or interacting with other transcription factors. We observed less GR in the nucleus of SphK1^-/-^ cells (Figure 7A-B), indicating that glucocorticoid signaling may be impaired by SphK1 deletion at a point upstream of *SphK1* transcription.

**Figure 7:**
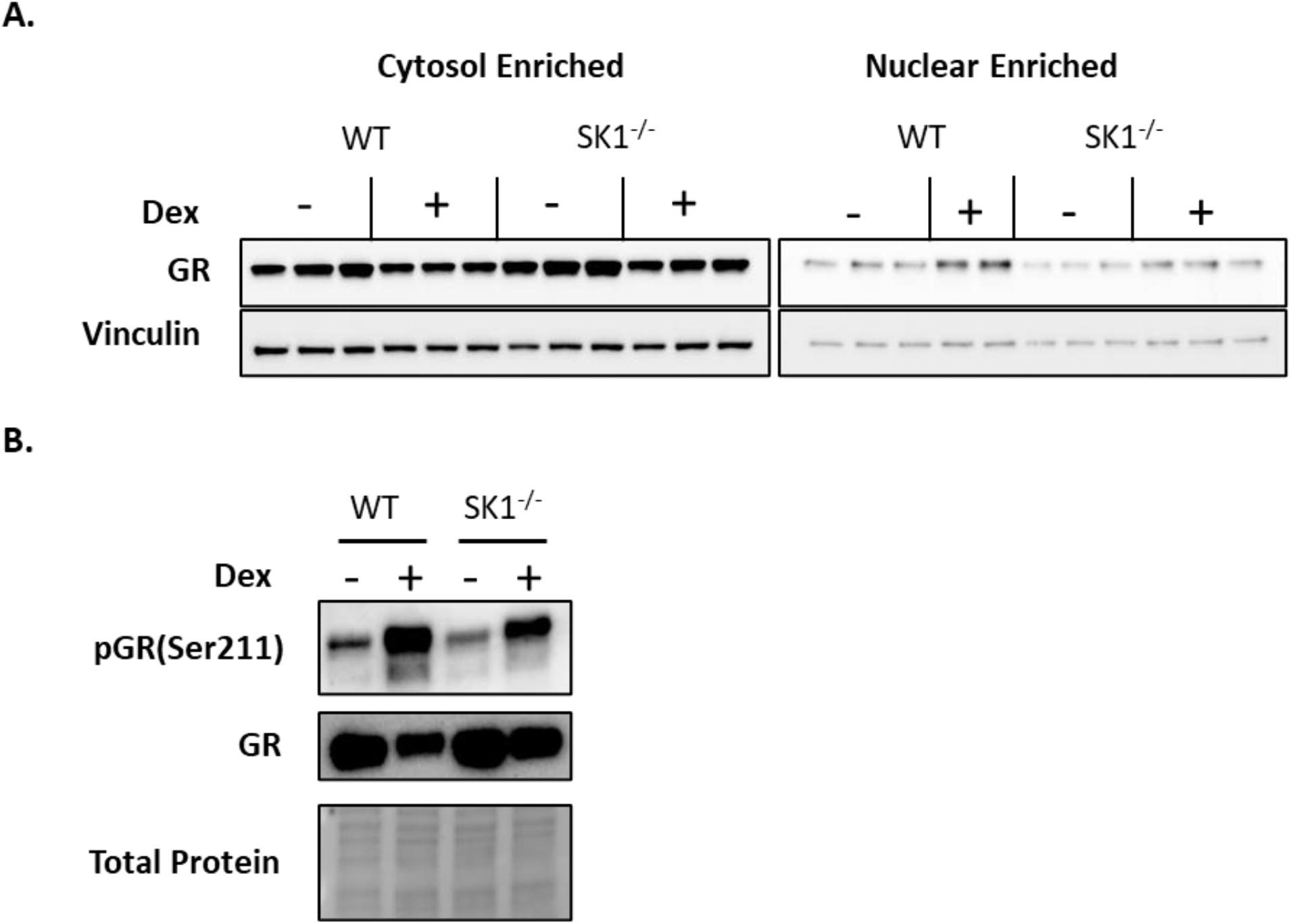
Glucocorticoid receptor activation in WT and SphK1^-/-^ ADSCs treated with dexamethasone. **A**. Glucocorticoid receptor translocation in WT and SphK1_-/-_ ADSCs stimulated with dexamethasone; Vinculin is a loading control. **B**. Phosphorylated and total GR in WT and SphK1_-/-_ whole cell lysates. n = 3. Due to the limitations of *in vitro* adipogenesis assays, we chose to investigate this further with *in vivo* experimentation. We treated SphK1^-/-^ mice and controls with corticosterone provided in drinking water for four weeks to determine if SphK1 mediates adipose tissue expansion as our *in vitro* data suggests. This form of chronic glucocorticoid exposure leads to robust adipose tissue expansion, redistribution, and insulin resistance, along with other metabolic effects such as liver steatosis and muscle wasting. Corticosterone-treated mice gained more weight than controls, consistent with other studies (Figure 8A-B)(24-26). Glucose tolerance tests were performed to determine if corticosterone impaired glucose handling differentially in SphK1^-/-^ mice. As previously reported, SphK1^-/-^ mice have reduced fasting glucose and AUC during glucose challenge (Figure 8C-D). Our data suggest this is not significantly affected by glucocorticoid treatment, though longer exposure to corticosterone may be needed to elicit measurable differences in glucose tolerance due to glucocorticoid exposure.

Next, the impact of corticosterone on adipose tissue mass was examined. Corticosterone led to a marked expansion of white adipose tissue in both SphK1^-/-^ and control mice. Gonadal and inguinal adipose depots were excised and weighed which revealed significantly lower expansion of these depots in SphK1^-/-^ mice (Figure 8E). Glucocorticoids have been associated with “whitening” of brown adipose tissue (25, 27). This was evident in corticosterone-treated mice, where it was difficult to distinguish clear boundaries between white and brown adipose tissue in Additionally, gene expression was measured in gonadal and inguinal adipose tissue homogenates to determine if genes related to adipogenesis were affected by corticosterone treatment and SphK1 deletion (Figure 8E). Corticosterone-treated control mice exhibited a ∼3-fold induction of SphK1 in gonadal but not inguinal adipose tissue (Figure 8F). This reveals a depot specific effect of corticosterone on adipose tissue. While not studied here, higher expression of glucocorticoid receptors in gonadal vs. subcutaneous adipose tissue has been demonstrated in rodents and humans which may pinpoint the depot specific effects of glucocorticoid action (28, 29). Expression of adipogenesis genes also varied by depot. Cebpd expression was reduced by corticosterone and significantly lower in SphK1^-/-^ in both the gonadal and inguinal depot. A similar trend was observed in the inguinal and gonadal depots from control mice, but this was not significant (p=0.18) (Figure 8G). This corresponds with our in vitro findings in ADSCs where Cebpd was lower in SphK1^-/-^ ADSCs. Alternatively, Cebpa was reduced in corticosterone treated mice of both genotypes in gonadal adipose tissue, but induced in the inguinal depot, and ∼2.5-fold higher in SphK1^-/-^ mice compared to controls (Figure 8H). Similarly, PPARγ, was reduced by corticosterone treatment in the gonadal depot of control mice and unchanged in SphK1^-/-^ mice, but significantly induced in the inguinal depot of SPHK1^-/-^ mice (Figure 8J). Sustained Cebpa and PPARγ expression are important for progression of adipogenesis, where Cebpd is an early trigger. Dlk1 (Pref-1) was basally higher in gonadal and inguinal depots of SphK1^-/-^ mice but reduced with corticosterone to a similar extent as controls (Figure 8I). These data highlight how systemic glucocorticoids act in a localized fashion, leading to different gene expression across adipose tissue depots.

**Figure 8:**
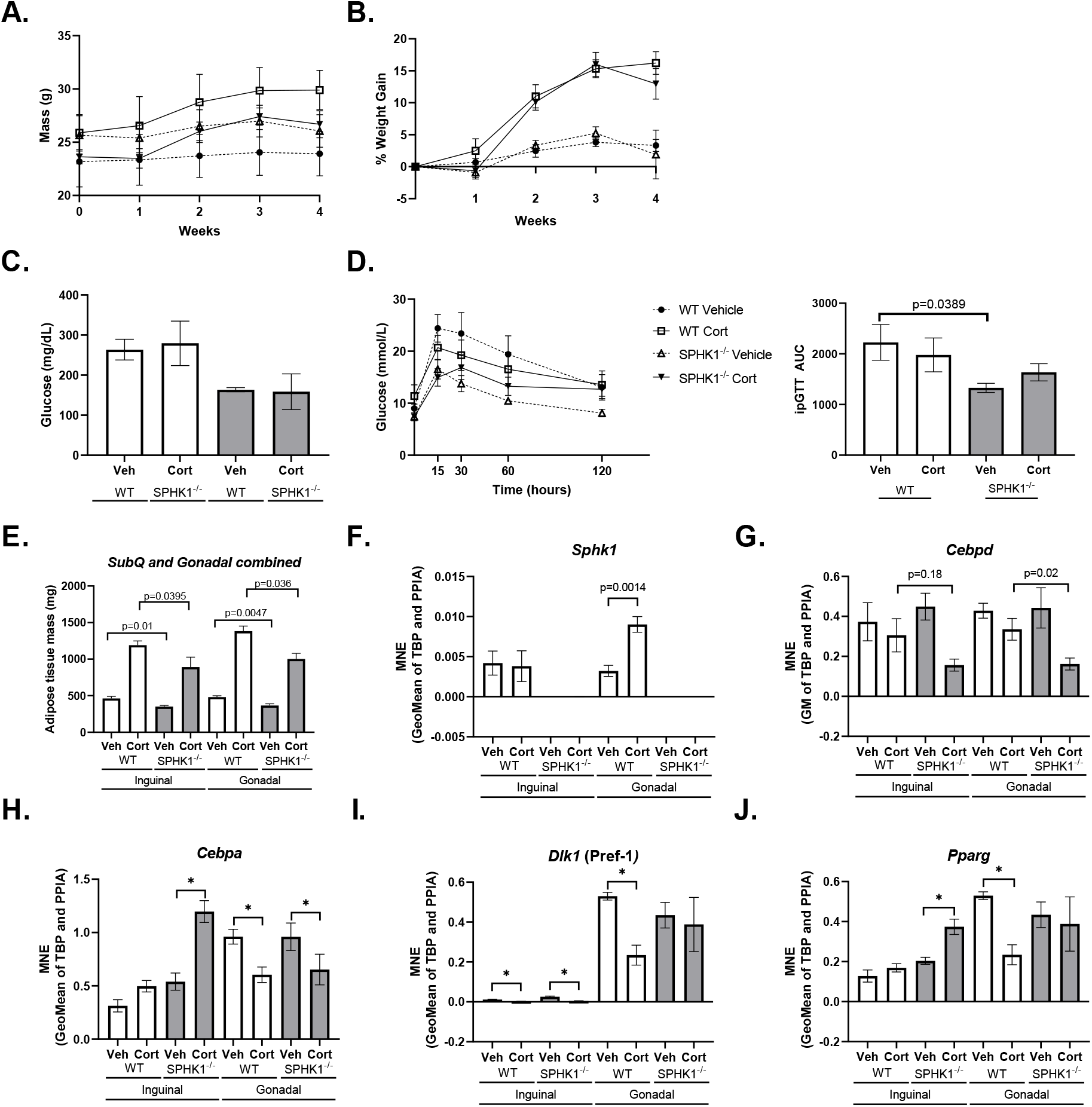
Corticosterone administration induces weight gain and has depot-specific effects on adipose tissue expansion that are altered in SphK1^-/-^ mice. **A**. Weight gain during corticosterone treatment. **B**. Percent weight gain during corticosterone treatment. **C**. Fasting glucose. **D**. Intraperitoneal glucose tolerance test and area under the curve (AUC). **E**. Inguinal and gonadal adipose tissue mass. **F-J**. Gene expression by qPCR in inguinal and gonadal adipose depots. Bars and dots represent mean ± SEM; n = 3. D. One way ANOVA. E.-J. unpaired t test. *P < 0.05. the interscapular region. Overall, more adipose tissue was found in this region, appearing beiger rather than distinct brown regions, though no clear differences were observed between genotypes (data not shown).

## Discussion

Our investigation into the role of SphK1 in adipogenesis were motivated by robust findings indicating adipocyte hypertrophy and reduced expression of mature adipocyte markers in adipose tissue of the SK1^fatKO^ mouse. Adipose hypertrophy due to impaired adipogenesis has previously been demonstrated because of adipose tissue dysfunction in obesity (30, 31). While we did not conduct detailed analysis of adipocyte cell number in the adipose tissue depots of SK1^fatKO^ mice and cannot definitively claim that adipocyte hypertrophy is the lone cause of their increased adipose tissue mass; we felt these findings were interesting enough to pursue potential functions of SphK1 in the adipogenic program. It is also relevant to note, that as SPHK1 deletion in SK1^fatKO^ mice is driven by adiponectin *Cre*, and adiponectin is only expressed in later stages of adipogenesis; any impacts of SphK1 deletion in this system are likely linked to later stage (adipocyte maturation) processes or the result cellular cross-talk with surrounding mature adipocytes lacking SphK1 within the tissue microenvironment.

In characterizing key timepoints of SphK1/S1P activation, we observed a robust induction of *SphK1* in early adipogenesis, which was attenuated through maturation, and is consistent with some published studies in 3T3-L1s (13, 32). Despite clear activation of SphK1 during adipogenesis, SphK1^-/-^ ADSCs were able to efficiently differentiate into functional adipocytes under standard adipogenesis conditions *in vitro*. This indicates that SphK1 is not critical for adipogenesis, at least when chemically induced, but may act as an enhancer of early adipogenic events. Interestingly, while PPARγ and FABP4 levels were reduced in SphK1^-/-^ cells upon induction, by 24 to 48 hours, they had surpassed those of control cells without any meaningful impacts on triglyceride accumulation as the cells matured (Figure 3). These markers must reach a threshold for adipogenesis to proceed, which indicates despite seemingly less committed than control ADSCs, SphK1^-/-^ cells are able to differentiate normally *in vitro*. These findings also bring into question if expression of SphK1 in later stages of adipogenesis may be inhibitory, allowing SphK1^-/-^ cells to “catch-up” to SphK1 expressing cells. Further investigation will be necessary to confirm any stage dependent effects of SphK1 on adipogenesis. The *in vitro* data presented here paints a similar picture to what was observed in SK1^fatKO^ mice fed high fat diet. Despite reduced expression of PPARγ and FABP4 in these mice, adipose expansion was able to occur, albeit seemingly more through hypertrophy than hyperplasia.

Treatment of ADSCs with individual pro-adipogenic media components revealed that SphK1 induction was entirely dependent on dexamethasone, suggesting a link between glucocorticoid and SphK1 activation. Indeed, sites of glucocorticoid receptor binding (glucocorticoid response elements) have been reported near the SphK1 promoter in a ChIP sequencing study of dexamethasone treated 3T3-L1 adipocytes (33). Furthermore, there have been reports of SphK1 induction by glucocorticoids in other cell types, including renal mesangial cells (34), macrophages (35) and human fibroblasts (36). These studies report a variety of outcomes downstream of SphK1, including prevention of apoptosis and inflammation. To our knowledge this is the first study to investigate the role of SphK1 in glucocorticoid signaling the adipocyte. Interestingly, treating with the full induction cocktail (insulin, IBMX, dexamethasone, rosiglitazone) induced *SphK1*, but to a much lower extent than dexamethasone alone (Figure 5A). This indicates that one of the other components may inhibit *SphK1* transcription, or the initiation of adipogenesis itself may limit *SphK1* expression. Additionally, as *Sphk2* expression was very low in ADSCs and was not responsive to dexamethasone treatment, we hypothesize that there is a function specific to SphK1 in the regulation of glucocorticoid signaling.

Dexamethasone induced expression of C/EBPδ and C/EBPα was lower in SphK1^-/-^ ADSCS treated with dexamethasone. Adipogenesis is orchestrated through a complex transcription factor network including C/EBPs to promote expression and maintenance of PPARγ, the “master regulator” (37, 38) Determining how SphK1/S1P interact with these key players is essential to understanding their role in adipogenesis.

As S1P receptor mediated signaling is often thought to be the primary mode of signaling downstream of SphK1, we were surprised to see no effect of S1P or receptor agonists on C/EBPs or adipogenesis. Moreover, we observed reductions in expression of S1P receptors upon dexamethasone treatment, further suggesting that S1P receptor mediated signaling is not occurring in our system. This is especially perplexing as several other groups have demonstrated negative effects of S1P receptor knockout or inhibition on adipogenesis (14-16). We hypothesize that the main effects of SphK1 in adipogenesis, at least in the context of glucocorticoid signaling are not receptor mediated. Intracellular mechanisms of SphK1/S1P will be of interest for future experiments.

Perhaps even more puzzling, is reduced GR in the nucleus, and reduced phosphorylated GR (Ser211) in whole-cell lysates from SphK1^-/-^ ADSCs treated with dexamethasone compared to controls. Both readouts are indicative of active glucocorticoid signaling, suggesting an impairment in SphK1^-/-^ cells. Dexamethasone stimulated SphK1 expression is expected to occur downstream of glucocorticoid receptor activation which suggests a positive feedback regulation of glucocorticoid receptor activation by SphK1/S1P.

Adipose tissue distribution is an important factor in overall metabolic health. Glucocorticoid administration to mice *in vivo* led to activation of *SphK1* in the gonadal adipose depot along with significant expansion of all adipose depots. This also occurred in SphK1^-/-^ mice but was significantly reduced. The depot specific activation of *SphK1* observed here reveals variations in SPHK1-dependent gene expression with respect to depot, a novel role for SphK1. These data suggest that SPHK1 deletion affects glucocorticoid-induced adipose tissue expansion directly but may also impact expansion of adipose depots in an indirect manner. SphK1 crosstalk between adipose tissue depots and other organs may explain some of the differences in gene expression we observed in the inguinal depot of SphK1^-/-^ mice treated with corticosterone.

One of our other initial findings, lower PDGFRβ expressing cells in SphK1^-/-^ mice provided additional evidence for a role of SphK1 in adipogenesis. PDGFRβ is one marker associated with adipocyte commitment (39). It is also known to activate SphK1 and promotes its translocation to the plasma membrane to stimulate signaling pathways involved in proliferation, detachment and migration (40). PDGFRβ has also been implicated in neovascularization during adipose tissue expansion (41). We propose that SphK1 promotes maintenance of PDGFRβ+ cell populations that can undergo adipogenesis when adipose tissue expansion is necessary. To our knowledge, no links between PDGFRβ and glucocorticoid signaling in the adipocyte have been demonstrated in existing literature. Though SphK1/S1P signaling in the context of PDGFRβ were not addressed in this study which focused on glucocorticoid mediated SphK1 signaling, there is potential SphK1 has additional roles independent of glucocorticoid signaling in the maintenance of pre-adipocyte populations within adipose tissue.

Overall, our results suggest that SphK1 is not strictly required for adipogenesis, but acts as an enhancer, particularly in the contexts of obesity and glucocorticoid excess. A similar conclusion was reached in an *in vivo* study showing that mice could still generate adipose tissue, despite compromised glucocorticoid signaling (42).Glucocorticoids paradoxically regulate both anabolic and catabolic pathways in adipocytes. Chronic glucocorticoid exposure causes a shift toward a positive energy balance, where anabolic effects exceed catabolism, leading to increased adipogenesis and overall adipose tissue expansion (33, 43). Though glucocorticoids affect many genes and signaling pathways, our data demonstrates that key adipogenesis genes are both glucocorticoid-responsive and downstream of SphK1 signaling. Consistently, *Cebpd* activation is reduced in SphK1^-/-^ cells in response to both acute glucocorticoid exposure in primary ADSCs and chronic exposure in mice. In summary, our findings support multiple roles for adipocyte-derived SphK1/S1P in adipogenesis, affecting whole-body physiology and driven by temporal and context-dependent factors. These results are summarized in Figure 9.

**Figure 9:**
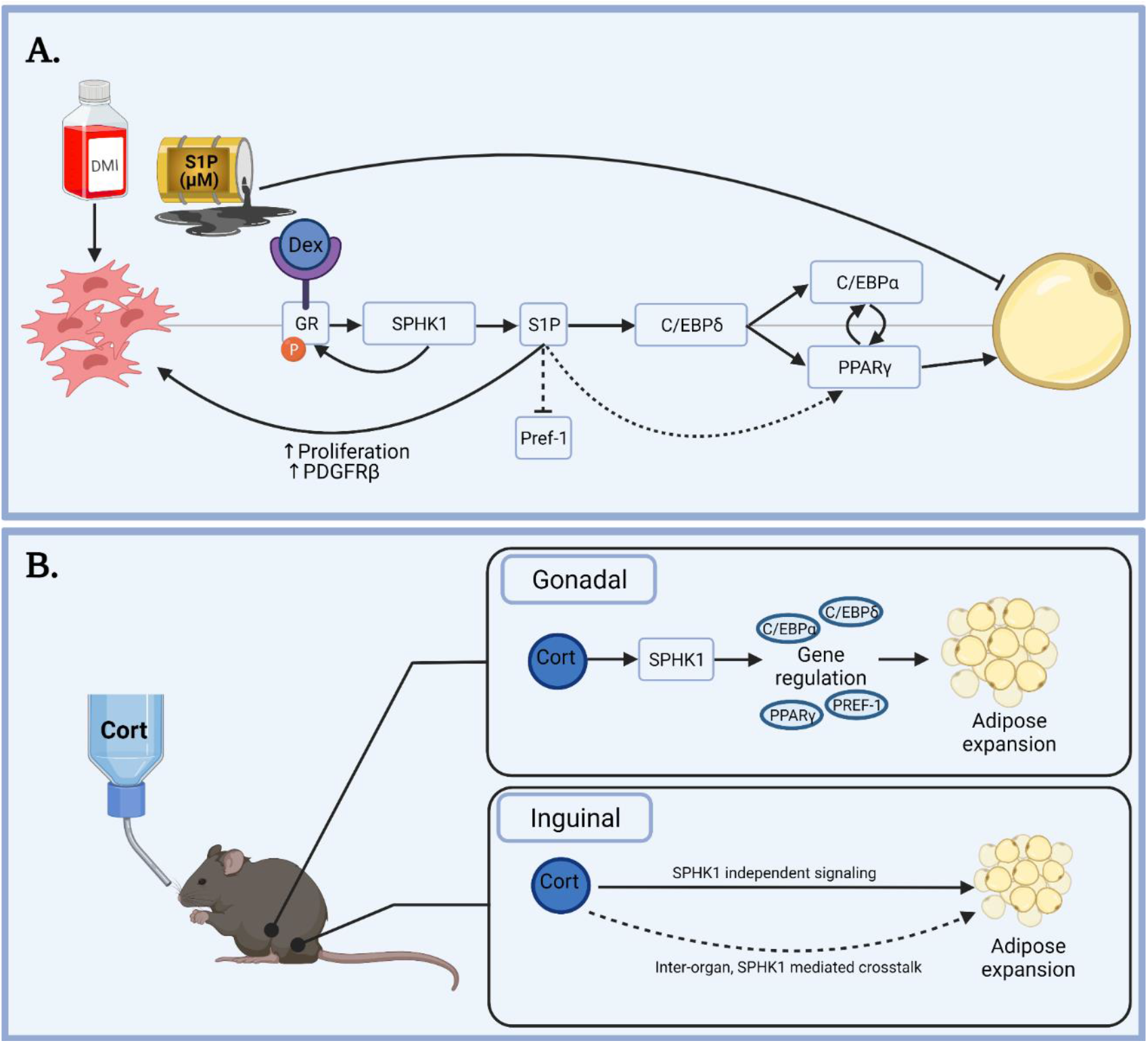
Summary of known and proposed roles of SPHK1/S1P in adipogenesis and adipose tissue depots. Solid lines represent mechanisms directly supported by our data or established pathways. Dashed lines represent proposed mechanisms or have been investigated in other cell types or contexts. S1P in micromolar concentrations has been shown to inhibit adipogenesis by several groups. However, SPHK1 expression and activity are induced early in adipogenesis in ADSCs treated with dexamethasone, IBMX, and insulin (DMI) with or without potent PPARγ agonist rosiglitazone. Dexamethasone (Dex) is the only component of adipogenic induction medium that induces SPHK1, indicating a role for SPHK1 in glucocorticoid signaling during adipogenesis. We observed reduced C/EBPδ activation after dexamethasone treatment, and reduced activated glucocorticoid receptor (GR) in SPHK1-/- cells. Potentially independent of glucocorticoid signaling, we also observed increased Pref-1, an inhibitor of adipogenesis, and reduced PDGFRβ. While not investigated in our system, S1P has previously been identified as a PPARγ agonist, another potential mechanism of SPHK1 in adipogenesis. Treatment of mice with corticosterone (Cort) in drinking water leads to SPHK1 dependent expression of adipogenesis genes and expansion of gonadal adipose tissue. SPHK1 is not induced in inguinal adipose tissue of mice administered corticosterone. Glucorcorticoid mediated expansion of this depot likely occurs via SPHK1 independent signaling events and possibly inter-organ SPHK1 mediated crosstalk.

## Acknowledgements

This study and its personnel were supported in part by grants to Dr Cowart from the National Institutes of Health (NIH; R01HL117233 and R01HL151243) and Veterans’ Affairs (IKBX006315 and 5I01BX000200). Grants F31HL156529 and T32HL149645 were provided to Dr Kovilakath; R21AA030647 to Drs Cowart and Montefusco. Services and products in support of the research project were generated by the following Virginia Commonwealth University/Massey Comprehensive Cancer Center Shared Resources: The Lipidomics and Metabolomics Shared Resource, the Cancer Mouse Models Core, the Microscopy Shared Resource, and the Transgenic/Knockout Mouse Shared Resource, supported in part with funding from NIH-NCI Cancer Center Support grant P30 CA016059.

